# Flexible analysis of TSS mapping data and detection of TSS shifts with TSRexploreR

**DOI:** 10.1101/2021.02.14.431114

**Authors:** Robert A. Policastro, Daniel J. McDonald, Volker P. Brendel, Gabriel E. Zentner

## Abstract

Heterogeneity in transcription initiation has important consequences for transcript stability and translation, and shifts in transcription start site (TSS) usage are prevalent in various disease and developmental contexts. Accordingly, numerous methods for global TSS profiling have been developed, including our recently published Survey of TRanscription Initiation at Promoter Elements with high-throughput sequencing (STRIPE-seq), a method to profile transcription start sites (TSSs) on a genome-wide scale with minimal cost and time. In parallel to our development of STRIPE-seq, we built TSRexploreR, an R package for end-to-end analysis of TSS mapping data. TSRexploreR provides functions for TSS and TSR detection, normalization, correlation, visualization, and differential TSS/TSR analysis. TSRexploreR is highly interoperable, accepting the data structures of TSS and TSR sets generated by several existing tools for processing and alignment of TSS mapping data, such as CAGEr for Cap Analysis of Gene Expression (CAGE) data. Lastly, TSRexploreR implements a novel approach for the detection of shifts in TSS distribution.

## Introduction

Genome-wide mapping of transcription start sites (TSSs) is crucial to understanding gene regulation. Clusters of TSSs, referred to as transcription start regions (TSRs), are associated with promoter elements and represent genomic positions at which RNA polymerase can initiate synthesis of new RNA molecules. Variation in TSS usage alters the length of 5’ untranslated regions (5’ UTRs), which has been shown to influence transcript stability and translation (1–3), and is recognized as a major contributor to transcript isoform diversity in mammalian cells and tissues (4–6). Alternative initiation has also been described in human cancers (7,8) and inflammatory bowel diseases (9) as well as during development, particularly in zebrafish (10). Thus, understanding gene regulation under physiological and pathologic conditions on a global scale requires accurate profiling of TSSs. To this end, numerous techniques have been developed, including Cap Analysis of Gene Expression (CAGE) (11), RNA Annotation and Mapping of Promoters for Analysis of Gene Expression (RAMPAGE) (12), and 5’ global run-on sequencing (GRO-cap) (13).

We recently introduced a new TSS mapping method termed Survey of TRanscription Initiation at Promoter Elements with high-throughput sequencing (STRIPE-seq) (14), a rapid, efficient, simple, and cost-effective TSS profiling approach compatible with limited input amounts. In parallel, we developed software to streamline analysis of STRIPE-seq data, as well as data resulting from other popular methods. Here, we describe the product of our code development as TSRexploreR, an R package for comprehensive and flexible exploration of TSS mapping data. TSRexploreR accepts pre-processed TSS and TSR data in a variety of common formats, including prior alignment results in BAM format. TSRexploreR performs normalization and offers a plethora of functions for correlation, visualization, and differential TSS/TSR analysis. Furthermore, TSRexploreR implements a novel approach to detect shifts in TSS distributions within TSRs. In sum, TSRexploreR is a feature-rich, interoperable, and easy-to-use software package for comprehensive analysis of TSS mapping data.

## Materials and Methods

### TSRexploreR implementation

TSRexploreR is fully implemented in R (with the exception of TSS shifting analysis, described below) and makes use of numerous Bioconductor packages and CRAN libraries such as Tidyverse and data.table. Data is stored in a TSRexploreR S4 object in common Bioconductor formats such as GenomicRanges (GRanges) or as a data.table for rapid, memory-efficient manipulation. TSRexplorer accepts bedGraph, bigWig, CTSS, and tab-delimited table files for TSSs and BED and tab-delimited table files for TSRs. Alignment BAM files can also be processed by TSRexploreR, as described below. TSRexploreR is packaged with STRIPE-seq-detected TSSs along budding yeast chromosome IV alongside the Ensembl release 99 budding yeast V64-1-1 genome sequence and annotation GTF. TSRexploreR is available at https://github.com/zentnerlab/TSRexploreR/releases/tag/v0.1.0 and as a Singularity container (library://zentlab/default/tsrexplorer:main), ensuring prolonged compatibility and reproducibility.

### BAM processing

Alignments in BAM format are loaded into TSRexploreR using the GenomicAlignments package (15) and can be processed as needed during import. An analysis of soft-clipping is performed, with reads having more than a user-specified number of soft-clipped bases removed. Filtering based on BAM flags is also performed, enabling removal of secondary alignments and, for paired-end reads, removal of unpaired or improperly paired reads and read pairs flagged as duplicates based on identical start and end positions. Overlapping 5’ read ends are then aggregated into TSSs. It has been frequently observed in both CAGE and TSRT-based TSS mapping protocols that a nonspecific G (corresponding to C on the first-strand cDNA) is often present at the 5’-most position of the R1 read (16,17). To correct for this artifact, we determine the frequency of reads with soft-clipped G bases. For each read with a 5’ G following removal of soft-clipped bases, a Bernoulli trial is conducted using the aforementioned soft-clipped G frequency as the “success” probability to decide if the G should be removed, which is similar in principle to the approach used with CAGE data (16).

### TSRexplorer vignettes

Step-by-step vignettes for performing common tasks in TSRexploreR are available at the following URLs:

BAM import and processing: https://github.com/zentnerlab/TSRexploreR/blob/v0.1.0/documentation/BAM_PROCESSING.pdf

Standard TSS/TSR exploration: https://github.com/zentnerlab/TSRexploreR/blob/v0.1.0/documentation/STANDARD_ANALYSIS.pdf

Differential feature analysis: https://github.com/zentnerlab/TSRexploreR/blob/v0.1.0/documentation/DIFF_FEATURES.pdf

TSS shifting analysis: https://github.com/zentnerlab/TSRexploreR/blob/v0.1.0/documentation/FEATURE_SHIFT.pdf

Data conditioning: https://github.com/zentnerlab/TSRexploreR/blob/v0.1.0/documentation/DATA_CONDITIONING.pdf

### TSS and TSR analysis

Yeast nAnT-iCAGE CTSSs (18) were obtained from www.yeastss.org (19) and imported into TSRexploreR. The nine growth conditions analyzed were: log-phase growth in rich yeast-peptone-dextrose medium (YPD, the control condition), cell cycle arrest with □-factor, DNA damage induced by methyl methanesulfonate (DD), diauxic shift (DSA), YP medium with 16% glucose to induce fermentation (Glc), log-phase growth in yeast-peptone-galactose medium (Gal), oxidative stress induced by H_2_O_2_, 37°C heat shock (HS), and osmotic stress induced by NaCl. For genome assembly and annotation we used the R packages ‘BSgenome.Scerevisiae.UCSC.sacCer3’ v1.4.0 and ‘TxDb.Scerevisiae.UCSC.sacCer3.sgdGene’ v3.2.2, respectively. Code used for yeast CAGE analysis is available at https://github.com/zentnerlab/Policastro_etal_2021/tree/v0.1.0.

### Zebrafish TSS shifting analysis

For TSRexploreR analysis, zebrafish developmental CAGE data were obtained as TPM-normalized bigWig files from http://promshift.genereg.net/zebrafish/CAGE/ and imported into TSRexploreR. Scores for negative-strand TSSs were multiplied by −1 to yield positive values. For CAGEr analysis, datasets were imported into CAGEr v1.30.3 from the R package ‘ZebrafishDevelopmentalCAGE’ v0.99.0 and TPM-normalized using the power-law approach. For both methods, TSSs supported by ≥3 TPM in one of the two samples were clustered into TSRs with a maximum clustering distance of 25 bp and a maximum TSR width of 250 bp. TSRs supported by at least 10 TPM in both samples were merged if they were within 100 bp of one another. An FDR threshold of 0.05 was used to assess the significance of shifting results from both approaches. Code used for shifting analysis is available at https://github.com/zentnerlab/Policastro_etal_2021/tree/v0.1.0.

## Results and Discussion

### Analysis of yeast nAnT-iCAGE data with TSRexploreR

To demonstrate the features of TSRexploreR, we analyzed nAnT-iCAGE CTSSs from yeast cells grown under a variety of conditions (18). In cases where a single plot is shown, this indicates a result from one YPD control replicate.

#### Genomic annotation and threshold exploration

Using annotations provided in GTF or TxDb format, TSRexploreR links TSSs to known genomic features (20). Assignment of TSSs to such features, particularly promoters, is useful in establishing a read threshold for downstream analyses. TSRexploreR threshold analysis determines the fraction of TSSs that is promoter-proximal and the number of transcripts or genes with at least one unique TSS across a range of raw read count thresholds. This analysis allows selection of a threshold that balances removal of likely artifacts (particularly weak TSSs in gene bodies) with detection of lowly abundant but legitimate promoter-proximal TSSs. Using a promoter definition of −250 to +100 bp relative to annotated gene starts (start codons for mRNA genes, TSSs for ncRNA genes), we selected a threshold of 10 counts/TSS, yielding promoter-proximal TSS fractions of 0.603-0.750 (Figure 1B, Supplementary Figure S1). Following annotation and thresholding, the distribution of TSSs relative to known genomic features can be visualized as stacked barplots (Figure 1C). A feature detection plot, wherein the number of genes or transcripts with at least one unique TSS position meeting the specified threshold is displayed, can also be generated (Figure 1D).

**Figure 1.**
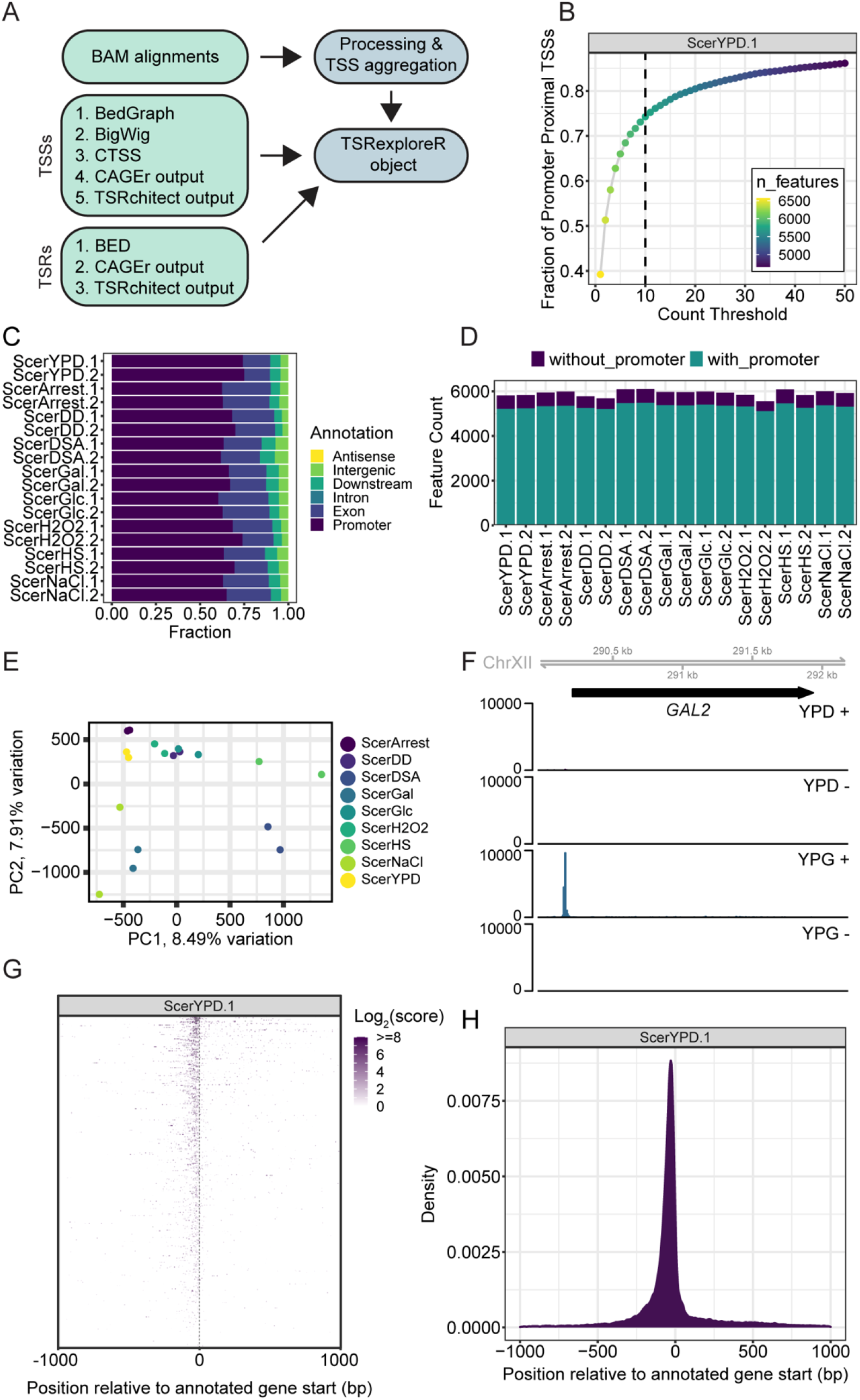
TSS analysis with TSRexploreR. (A) Schematic depicting input formats accepted by TSRexploreR and creation of the TSRexploreR object. (B) Threshold plot showing the fraction of TSSs that is promoter-proximal (−250 to +100 relative to annotated gene starts) and the number of features (in this case, transcripts) with at least one unique TSS position at each threshold in YPD replicate 1. (C) Barplot of the genomic distribution of TSSs in each sample. (D) Barplot of the number of transcripts with a unique TSS position in each sample, and whether that TSS is promoter-proximal or not. (E) PCA plot of TSSs detected in each CAGE sample. (F) Signal tracks of normalized signal (YPD and Gal replicate 1) at the *GAL2* locus. (G) Heatmap of normalized signal from YPD replicate 1 relative to annotated gene starts, sorted descending by total signal. (H) Density plot of unique TSS positions relative to annotated gene starts for YPD replicate 1.

#### Normalization

TSRexploreR includes three options for normalization. The first, counts per million (CPM), is a simple read number-based normalization approach commonly used for data visualization and is particularly appropriate for replicate comparison. However, CPM normalization is considered to be simplistic when comparing samples from distinct biological conditions (21) and so we implemented two additional normalization approaches considered more appropriate for such cases: trimmed mean of M-values (TMM) (21), used in edgeR (22), and median-of-ratios (MOR), used in DESeq2 (23). For this example, data were normalized using the MOR approach. Normalized samples can be compared via a PCA plot (24) (Figure 1E) or correlation heatmaps (25) (Supplementary Figure S2).

#### Visualization of TSS data

TSSs can be exported in bedGraph, bigWig, or tab-delimited table format. TSSs and/or TSRs at a specific gene or its promoter can also be directly visualized using Gviz (26). To demonstrate this feature, we plotted MOR-normalized TSS counts from one replicate each of the YPD and Gal conditions at the promoter of the *GAL2* gene, encoding a permease required for galactose utilization (Figure 1F). TSS signal around gene starts can also be displayed as a heatmap (Figure 1G), and the distribution of TSS positions relative to annotated gene starts can be visualized as a density plot (Figure 1H).

#### TSR detection and analysis

TSRexploreR uses a simple distance-based clustering approach to aggregate TSSs into TSRs based on a user-specified TSS count threshold that must be met in a specified number of samples and maximum inter-TSS distance. For this analysis, we used a raw count threshold of 10 in at least one sample, a maximum distance of 25 bp, and a maximum TSR width of 250. Many of the analyses described above for TSSs can also be applied to TSRs: correlation, analysis of genomic distribution, feature detection, and density/signal relative to annotated gene starts.

### Characterization of TSR features

It has been well established that there is a continuum of TSR shapes ranging from sharp or peaked, wherein transcription initiates at one or a few strong TSSs, to broad or dispersed, wherein there are several TSSs of similar strength (27). TSRexploreR calculates three metrics relating to TSR shape: 1) shape index (SI), which assesses the shape of TSRs via analysis of the position of each constituent TSS and its strength relative to the overall score of the TSR (27); 2) inter-quantile range (IQR), which measures the distance between the base positions of the given TSS signal quantiles, providing information on the width of a TSR without being affected by weak TSSs on its edges (28); 3) peak balance, which assesses the skew of TSSs around the TSR center (29) (Supplementary Figure S3).

### TSS sequence analysis

TSRexploreR includes several functions for analyzing the sequence context of TSSs. Furthermore, as it is often desirable to assess specific subsets of TSSs or TSRs, TSRexploreR includes a number of conditioning functions for grouping, ordering, quantiling, and filtering data. Here, we demonstrate these features by splitting TSRs into quintiles by score and plotting sequence logos (30) around the dominant TSS of each TSR in its quintile (Figure 2A). This analysis indicates a pyrimidine (Y = C or T) preference at the −1 position and a purine (R = A or G) preference at the +1 position (the TSS itself), as well as an A base in the −8 position (Figure 2A), consistent with previous studies (31), that decreased in information content with decreasing TSS score. Sequences surrounding TSSs can also be visualized as color plots (Figure 2B). Lastly, the frequencies of all observed −1/+1 dinucleotides can be plotted (Figure 2C).

**Figure 2.**
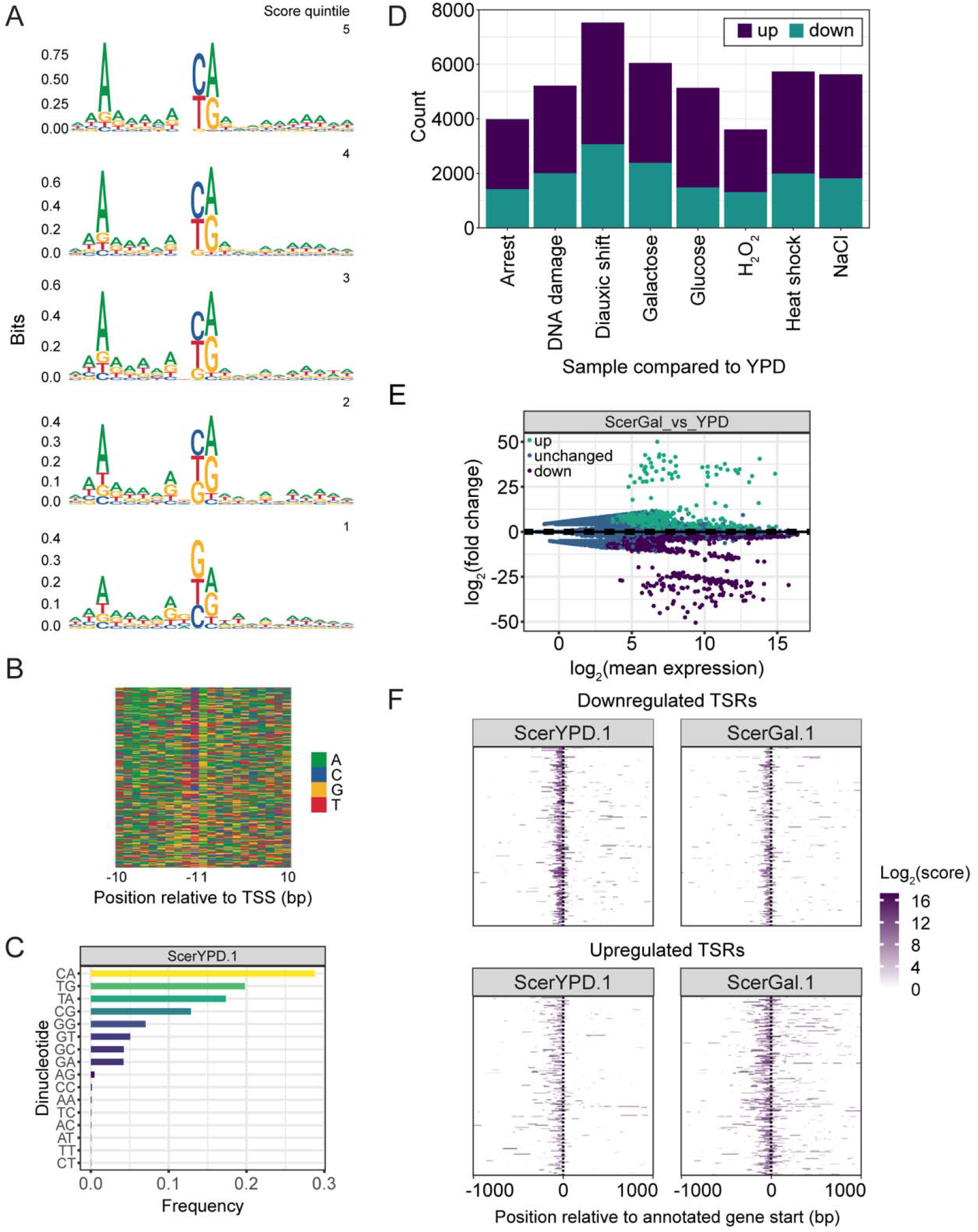
Sequence analysis and differential TSR detection. (A) Sequence logos (quintiled in descending order by TSR score), (B) base color plot (in descending order by TSR score), and (C) barplot of dinucleotide frequencies at the dominant TSSs of TSRs called in YPD replicate 1. Only TSRs > 10 bp in width were considered for these plots. (D) Barplot of the number of differentially expressed TSRs for comparison of each indicated condition to the YPD control. (E) MA plot of differential TSR results for the Gal versus YPD comparison. (F) Heatmap of normalized signal TSR around the annotated starts of genes with downregulated and upregulated promoter-proximal TSRs in the Gal versus YPD comparison.

### Analysis of differential TSR usage

Transcription is highly dynamic and plastic, able to respond quickly to myriad stimuli. To enable analysis of differential TSS and TSR usage across distinct conditions, TSRexploreR generates matrices of counts within merged regions that are used as input for edgeR or DESeq2. We used DESeq2 to build a statistical model and then performed contrasts of treated samples versus the control YPD condition (see Supplementary Table S1 for full differential TSR analysis results). As an overview of differential feature analysis, a stacked barplot of the number of changed features in each contrast can be generated (Figure 2D). The results of individual comparisons (for this example, Gal versus YPD) can also be visualized as a MA plot, displaying log_2_(fold change) versus mean expression (Figure 2E), and as a volcano plot, displaying -log_10_(FDR) versus log_2_(fold change) (Supplementary Figure S4). We also demonstrate visualization of differential TSR signal using conditioned heatmaps, wherein data are split by the direction of signal change and ordered by signal in the control sample (Figure 2F). To facilitate interpretation of differential feature analysis results, TSRexploreR annotates differential features and exports a list of gene names compatible with clusterProfiler, a robust R package for gene ontology (GO) analysis (32). Genes associated with upregulated promoter-proximal TSRs were enriched for GO biological process terms including ‘carbohydrate metabolic process’ and ‘generation of precursor metabolites and energy’ (Supplementary Figure S5, Supplementary Table S2). Genes with downregulated promoter-proximal TSRs were enriched for processes related to ribosome biogenesis (Supplementary Figure S5, Supplementary Table S2), consistent with previous work showing reduced levels of ribosomal protein gene transcripts in cells grown continuously in galactose (33).

### Detection of TSS shifts with TSRexploreR

Numerous studies indicate that large-scale shifts in TSS distribution are prevalent in various developmental settings (10) and may also be induced by mutations in general transcription factors (34). Computational detection of TSS shifts may be approached as testing for differences between two discrete probability distributions. The CAGEr package (28) assesses spatial shifts in TSS usage by generating aggregate TSRs from TSRs identified in all samples, and comparing empirical cumulative distribution functions (ECDFs), where the sample with larger total signal has its ECDF rescaled by the ratio of total signal. This results in a score between negative infinity and 1, with larger positive values posited to indicate that a given proportion of signal in the second sample is outside of the TSS-containing region in the first sample. For instance, a CAGEr shift score of 0.4 would indicate that at least 40% of the transcription initiation in the second sample is independent of that in the first sample. This approach only assesses spatial separation between two distributions and so does not address shifts in signal distribution at largely overlapping positions. Furthermore, it produces a substantial number of negative shift scores, the interpretation of which can be unclear. Lastly, it does not indicate shift direction. In addition to calculating the shifting score, CAGEr also performs a Kolmogorov-Smirnov (K-S) test on the ECDFs, identifying the point of maximal distance between them. The stated purpose of the K-S test is twofold: 1) to assess significance of the observed difference in TSS distribution between the two samples and 2) to capture changes in TSS distributions within mostly overlapping positions that are not captured by the shift score. However, calculation of the K-S statistic is unrelated to the shift score and so its p-value does not indicate the score’s significance. Furthermore, the derivation of the p-value formula for the K-S test assumes the data come from a continuous distribution, an assumption not met by TSS distributions, which are observed at discrete locations. Lastly, the K-S test, like the shift score, does not indicate direction.

Given these limitations, we implement an alternative approach to detecting TSS shifts using a more intuitive metric. We use a signed version of earth mover’s distance (EMD) (35), which we refer to as earth mover’s score (EMS), to characterize between-sample differences in TSS distributions within merged TSRs. For this approach, we imagine that the two TSS distributions in question are piles of dirt, and ask how much dirt from one pile we would need to move, how far, and in which direction, to recreate the distribution of the other sample. The computed EMS thus represents the minimum “cost” of converting one distribution into the other. The resulting score is between −1 and 1, with larger magnitudes indicating larger shifts and the sign indicating direction (negative values indicate upstream shifts and positive values indicate downstream shifts). Figure 3A-C illustrates the intuition for calculation of the EMS. Our implementation calculates shift scores as well as a p-value and FDR threshold based on a permutation test. TSS shifting analysis is implemented in C++ to enhance execution speed.

**Figure 3.**
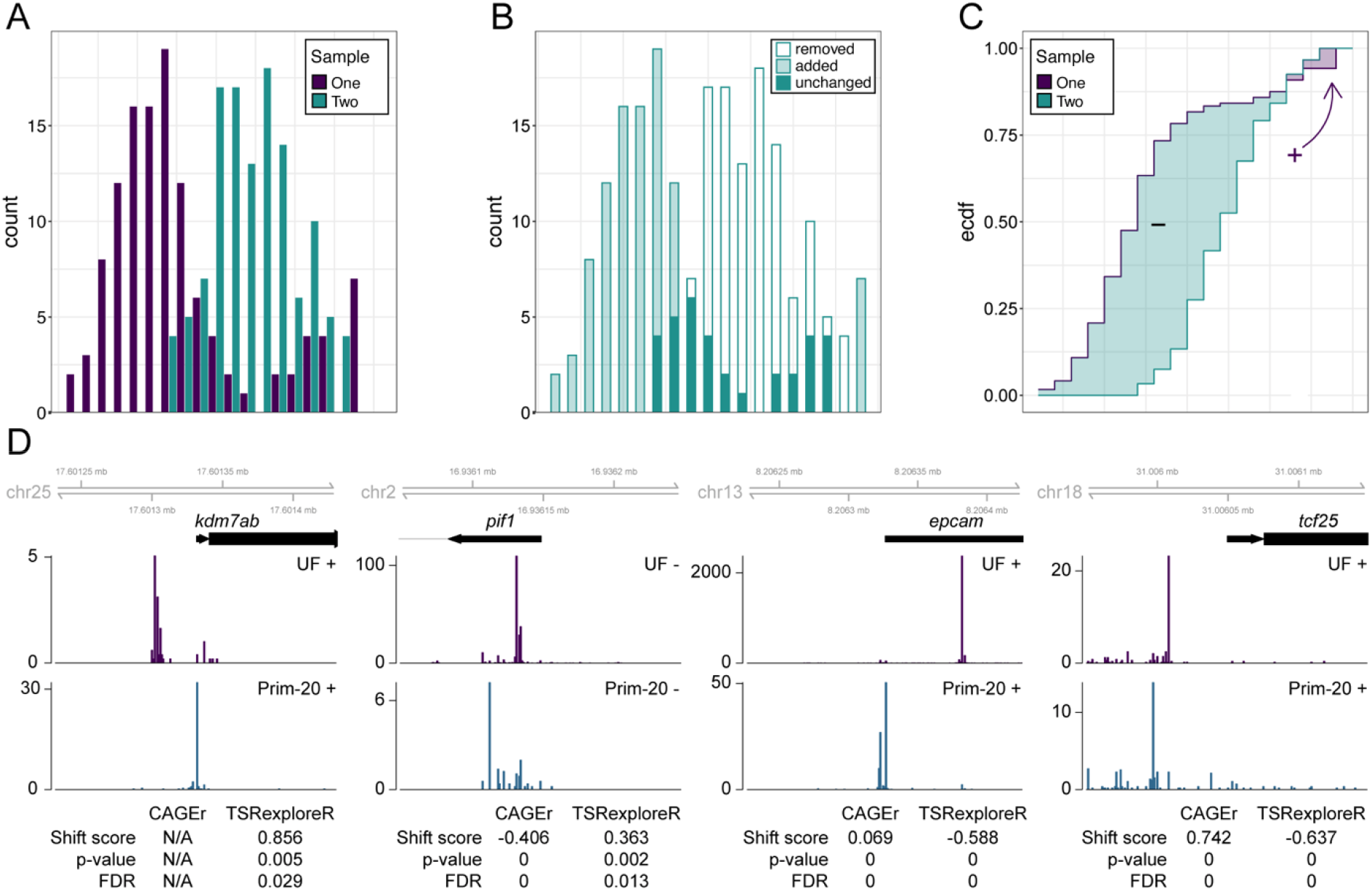
Detection of TSS shifts using earth mover’s score. (A) Stylized TSS distributions for two samples at a hypothetical region of interest. (B) Illustration of how Sample Two would need to be “moved” in order to match Sample One. Material (or “earth”) must be moved from the empty bars into the shaded bars while the solid bars remain unchanged. Some material has to be shifted in both directions, but more is moved upstream than downstream. Calculating how much, how far, and which “piles” to move is a standard constrained optimization problem known by the name “optimal transport”, but here it reduces to a simple integral. (C) Calculation of the EMS for the hypothetical example illustrated in (B). The upstream (purple, negative) and downstream (green, positive) areas between the ECDFs are simply integrated and then subtracted from each other. The result is normalized to the number of locations with expression in either sample. This example has an EMS of 0.243 with a p-value of 0 based on an approximate permutation test using 1000 resamples. (D) Tracks of zebrafish CAGE signal from the unfertilized egg (UF) and Prim-20 stages at four promoters displaying significant shifts in TSS distribution by the EMS-based method implemented in TSRexploreR. Note that only the strand from which the TSS signal originates is shown. Shift scores and statistical measures calculated by CAGEr and TSRexploreR are also included.

To illustrate the capacity of our EMS-based approach to detect TSS shifts, we turned to a set of CAGE experiments performed throughout zebrafish embryonic development. Detailed analysis of this dataset revealed distinct distributions of TSSs for the maternally-deposited and zygotic forms of several hundred transcripts (10), and so it provides a robust test case. We compared the earliest and latest time points assayed (unfertilized egg and Prim-20, respectively) using both the established CAGEr approach and our EMS-based method. Using CAGEr with no shift score threshold, we detected 3,950 significantly shifted TSRs, while applying a shift score threshold of 0.4 yielded 1,314 significant shifts (Supplementary Table S3). EMS-based analysis yielded 1,052 significantly shifted TSRs (Supplementary Table S4); we note that this number is slightly variable due to the use of a permutation test for determining significance. To illustrate the relationship between our EMS-based shift score and TSS redistribution, we visualized data at several loci displaying various degrees of shifting (Figure 3D). At *kdm7ab,* CAGE signal was markedly shifted downstream in the Prim-20 sample, yielded a shift score of 0.856 (FDR = 0.029); this shift was not detected by CAGEr. A more modest downstream shift was observed at *pif1* (shift score = 0.363, FDR = 0.013). The *pif1* shift was detected as highly significant by CAGEr (FDR = 0), though the shift score was negative (−0.406). At *epcam,* we detected a robust upstream shift (shift score = −0.588, FDR = 0); this shift was also detected by CAGEr, though with a very small shift score (0.069). Lastly, at *tcf25,* we observed a highly significant upstream shift (shift score = −0.637, FDR = 0). The *tcf25* shift was also marked as highly significant by CAGEr (FDR = 0), but the sign of the robust shift score (0.742) does not relate to the direction of the shift.

### Concluding remarks

TSRexploreR leverages the extensive Bioconductor ecosystem and the tidyverse to provide a feature-rich and straightforward tool for TSS mapping analysis. We also introduce a novel approach to detecting TSS shifts. While TSRexploreR was originally developed to handle STRIPE-seq data, it has been made highly interoperable and can thus be readily incorporated into workflows using existing TSS analysis software such as CAGEr, TSRchitect, and CAGEfightR.

## Acknowledgements

We thank Jason Tourigny for assistance with testing TSRexploreR functions.

## Funding

This work was supported by National Science Foundation grant IOS-1221984 [to V.P.B.] and National Institutes of Health grant R35GM128631 [to G.E.Z]

## Notes

https://github.com/zentnerlab/TSRexploreR/releases/tag/v0.1.0

### Competing Interest Statement

The authors have declared no competing interest.

