## Supplementary Material for "Flexible analysis of TSS mapping data and detection of TSS shifts with TSRexploreR"

Contents:
Supplementary Figures S1-S5
Legends for Supplementary Tables S1-S4

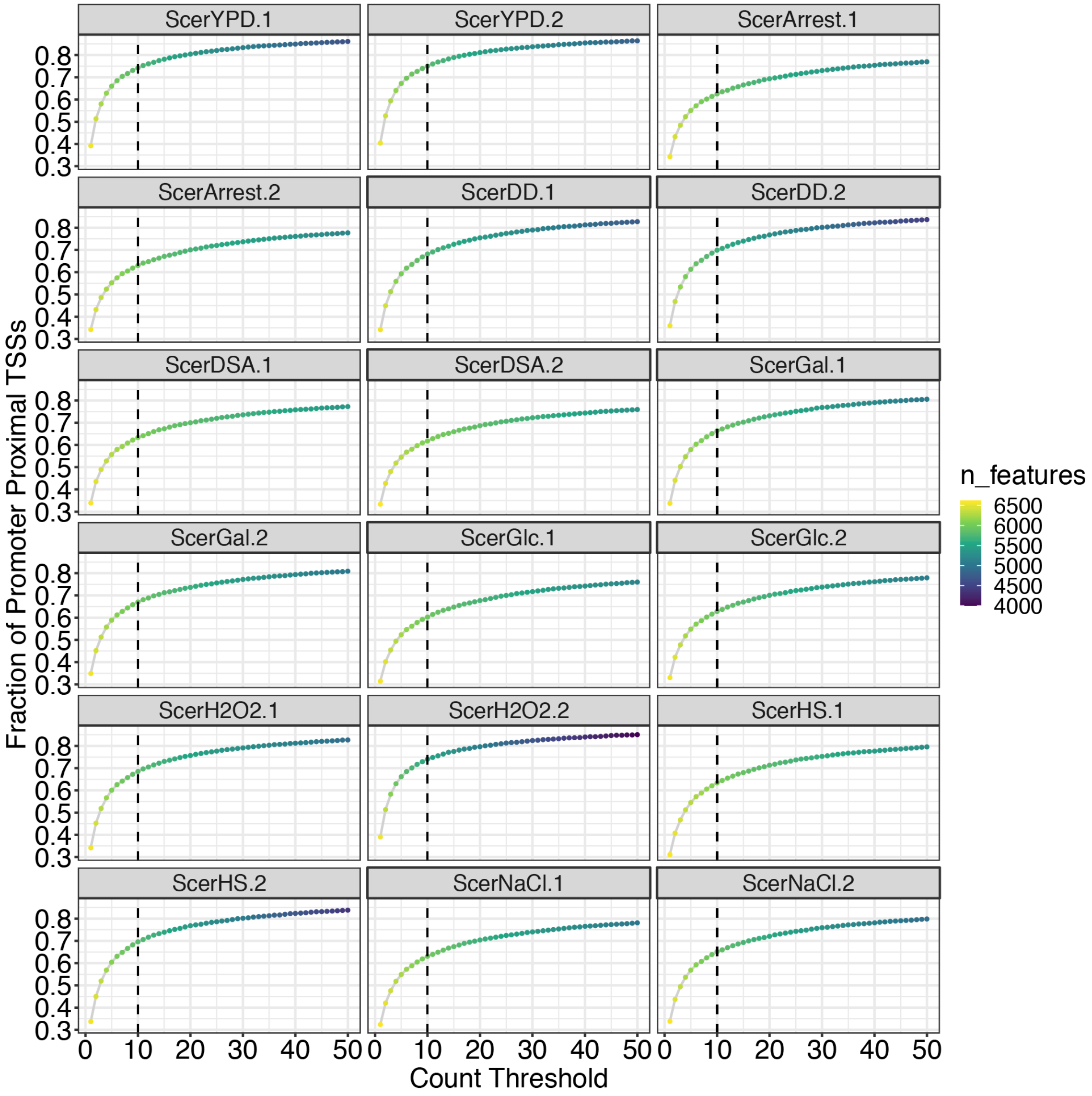


**Figure S1. Threshold plots for all yeast CAGE samples**

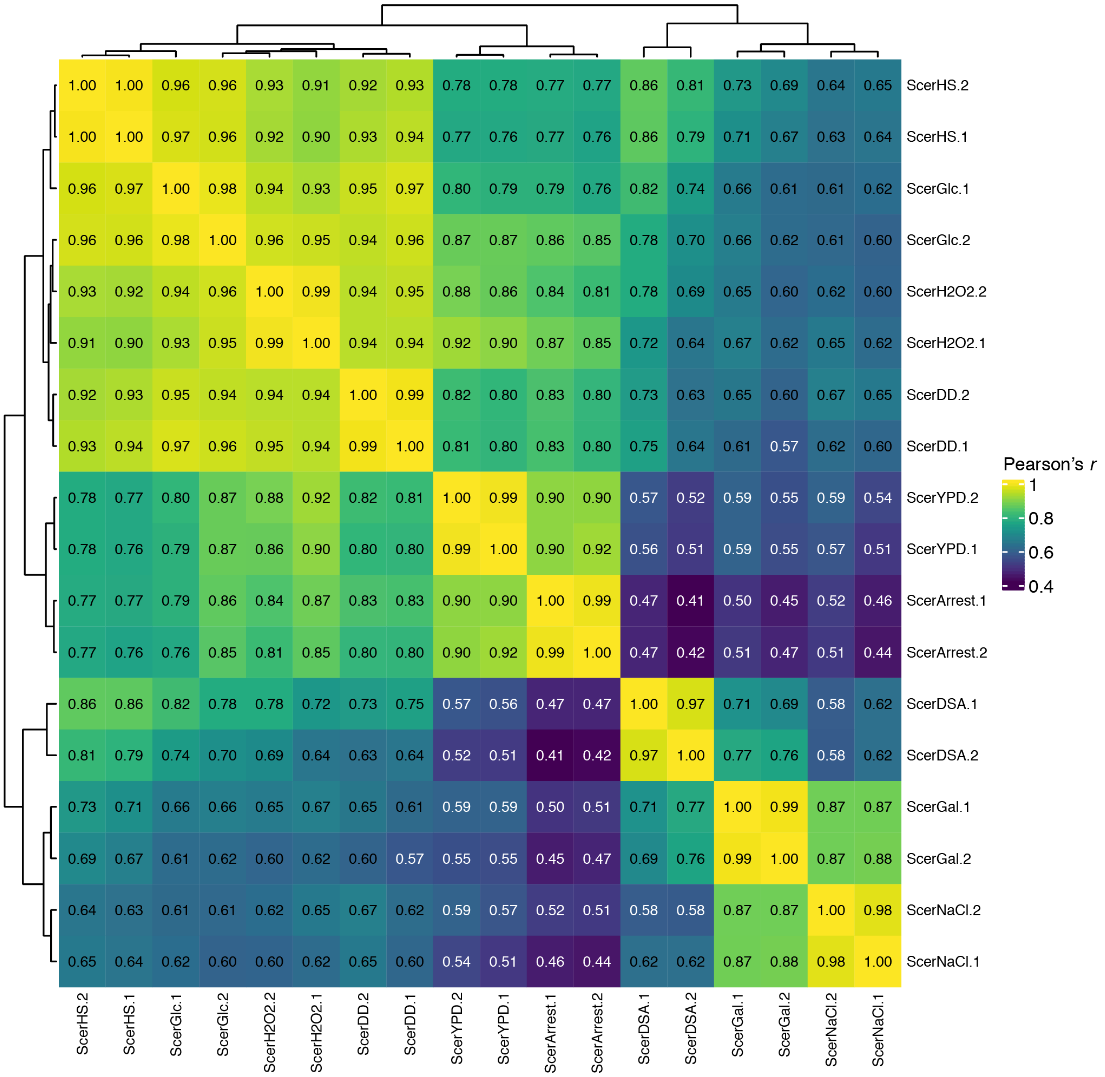


**Figure S2. Correlation analysis of yeast CAGE TSSs**

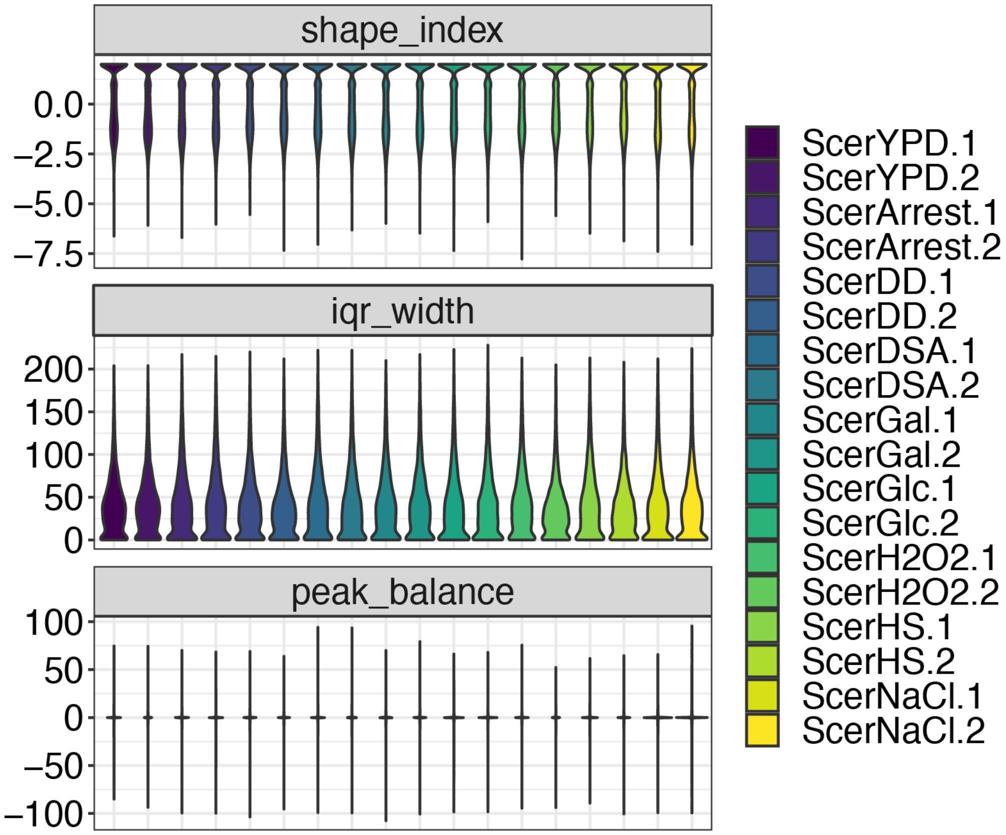


**Figure S3. Shape analysis of yeast CAGE TSRs**

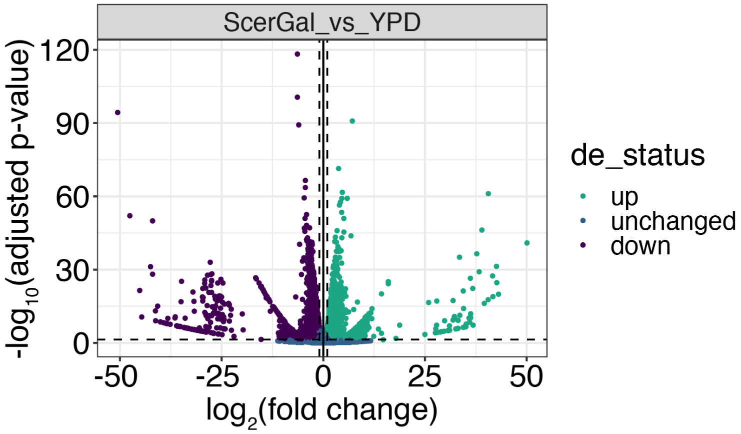


**Figure S4. Volcano plot of differential TSR analysis results of Gal versus YPD CAGE samples**

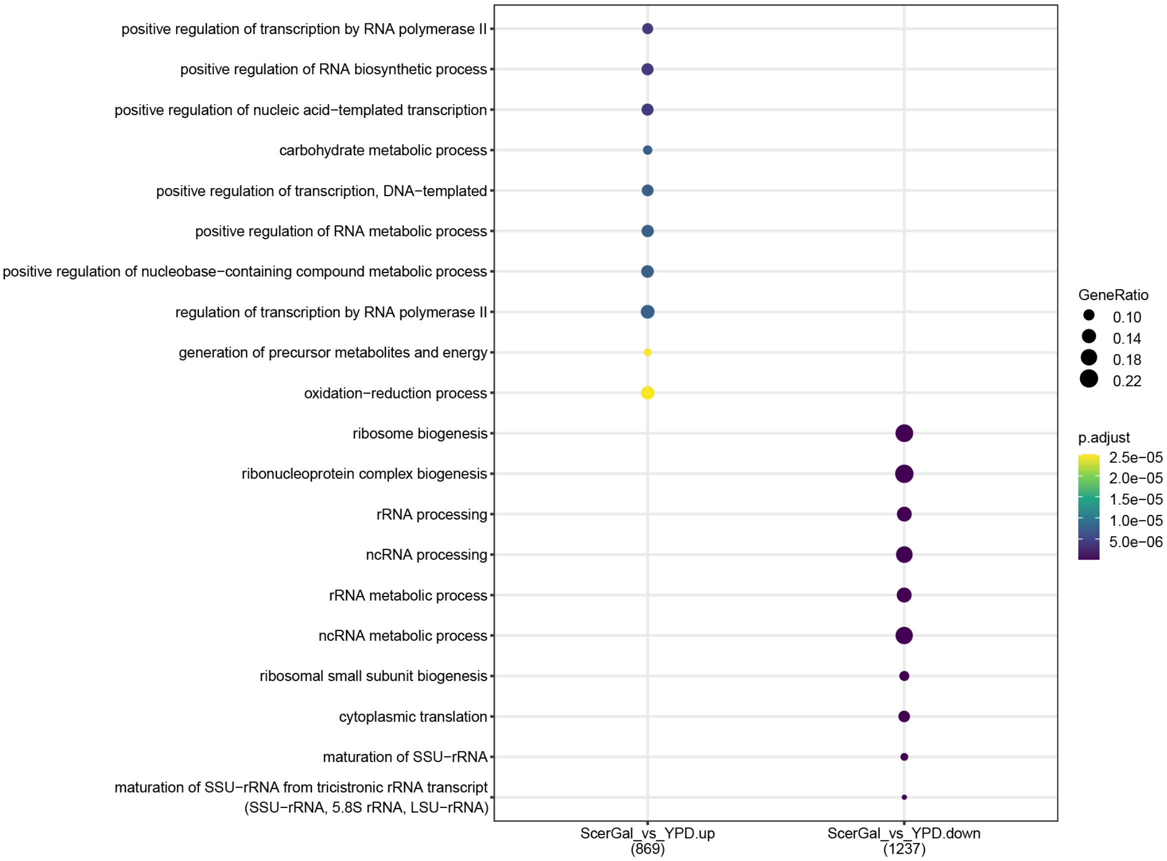


**Figure S5. GO Biological Process ontologies associated with upregulated and downregulated TSRs in the Gal versus YPD comparison**

**Supplementary Table legends**

**Table S1.** Full differential TSR analysis results.

**Table S2.** Full clusterProfiler GO biological process results for promoter-proximal differential TSRs in the Gal versus YPD comparison.

**Table S3.** CAGEr shifting analysis results.

**Table S4.** TSRexploreR shifting analysis results.
